# Highly reproducible murine model of oral *Salmonella* infection via inoculated food

**DOI:** 10.1101/593442

**Authors:** Olof R. Nilsson, Laszlo Kari, Olivia Steele-Mortimer

## Abstract

Oral infection of mice with *Salmonella* Typhimurium is an important model system. In particular C57Bl/6 mice, which are susceptible to *Salmonella*, are used to study both systemic and gastrointestinal pathogenesis. Pretreatment with streptomycin disrupts the intestinal microbiota and results in colitis resembling human intestinal *Salmonellosis*. Oral gavage is typically used for delivery of both antibiotic and bacteria. Although convenient, this method requires a moderate level of expertise, can be stressful for experimental animals, and may lead to unwanted tracheal or systemic introduction of bacteria. Here, we demonstrate a simple method for oral infection of mice using small pieces of regular mouse chow inoculated with a known number of bacteria. Mice readily ate chow pieces containing up to 10^8^ CFU *Salmonella*, allowing for a wide range of infectious doses. In mice pretreated with streptomycin, infection with inoculated chow resulted in less variability in numbers of bacteria recovered from tissues compared to oral gavage, and highly consistent infections even at doses as low as 10^3^ *Salmonella*. Mice not treated with streptomycin, as well as resistant Nramp1 reconstituted C57Bl/6J mice, were also readily infected using this method. In summary, we show that foodborne infection of mice by feeding with pieces of chow inoculated with *Salmonella* results in infection comparable to oral gavage but represents a natural route of infection with fewer side effects and less variability among mice.

## Introduction

Bacteria belonging to the genus *Salmonella enterica* subsp. *enterica* are one of the most common causes of self-limiting foodborne diarrheal disease in humans and other animals [1] and a leading cause of death due to foodborne pathogens globally [2] and in the US [3]. *Salmonella enterica* serovar Typhimurium (hereafter *Salmonella*) is one of the serovars most commonly isolated from human gastrointestinal infections. *Salmonella* infects new hosts in a fecal-to-oral manner and is often the cause of foodborne disease outbreaks when present in contaminated food, such as fresh produce [4] and poultry and egg products [5]. *Salmonella* has been thoroughly studied and is one of the best characterized human pathogens. This, combined with its simple growth requirements, has led to its frequent use as a model organism for *in vivo* studies of the pathogenesis of gastrointestinal infections.

The most widely used animal model for *Salmonella* is the mouse [6]. Strains of mice differ in their susceptibility to *Salmonella* infection, with C57Bl/6J and BALB/c mice being highly susceptible and 129/Sv strains of mice being very resistant [7–10]. Susceptibility is multifactorial, but one important resistance factor is the Nramp1 protein encoded by the *Slc11a1* gene [11]. Nramp1 is an ion transporter responsible for the transport of divalent cations out of phagosomes, thus limiting the availability of iron and other ions for ingested microbes and impairing their growth in phagocytes [12]. Many susceptible mouse strains, including C57Bl/6J, harbor a point mutation in the *Slc11a1* gene resulting in a non-functional Nramp1 protein [13, 14]. Oral infection of susceptible mouse strains eventually leads to a lethal systemic infection but without diarrhea and only diffuse enteritis [15]. However, following disruption of the intestinal microbiota by antibiotic treatment, mice develop intestinal inflammation more similar to human intestinal *Salmonellosis*, although *Salmonella* do still go systemic [16, 17].

In the preferred murine model of oral *Salmonella* infection, C57Bl/6 mice are treated with antibiotics and infected by oral gavage using blunt end gavage needles. This method of delivery is intragastric rather than oral, since substances are delivered directly into the stomach. While the use of gavage needles allows for the delivery of precise amounts of inoculum and timing of delivery, there are several drawbacks to their use, which have been recognized mostly in toxicological studies [18]. Performing oral gavage requires a moderate degree of technical expertise and can induce stress, e.g. raising corticosteroid levels in the blood or increasing blood pressure, which may affect study outcome [19–21]. Furthermore, mice may regurgitate delivered substances or infectious agents following gavage, resulting in tracheal or nasal administration [22, 23]. Lastly, gavage may induce pharyngeal or esophageal trauma, leading to the inadvertent delivery of substances or infectious agents directly into the blood stream or, in rare cases, death [23, 24].

Improvements to oral gavage have been suggested, such as precoating needles with sucrose, which improved gavage success rate and reduced stress of animals [25]. An alternative method to gavage would be ingestion of food or water containing a pathogen. This would circumvent many of the drawbacks with oral gavage and mimics a natural route of infection for *Salmonella*. Recently, a paper described a method of delivering *Salmonella* orally in drinking water [26], and a similar approach was used for the oral delivery of a *Salmonella* vaccine to sheep [27]. Inoculated food (pieces of bread) has been used for *Listeria monocytogenes* infections [28]. However, to our knowledge, food as a vehicle of delivery for *Salmonella* infection has not been reported.

In this paper, we describe an oral infection method using pieces of regular mouse chow inoculated with *Salmonella*. Preparation of inoculated chow is simple and mice readily consume chow containing high numbers of bacteria. This mode of infection leads to a very consistent disease progression among mice, less variability in bacterial load, in both systemic and gastrointestinal tissues, and eliminates many of the possible drawbacks with oral gavage. Importantly, this method represents a natural route of infection with *Salmonella*.

## Results and Discussion

### *Salmonella* remains viable on mouse chow

We hypothesized that regular mouse chow could be used to inoculate mice with *Salmonella* to establish a simple, stress-free and natural route of *Salmonella* infection. At our facility, mice typically receive 2016 Teklad Global 16% Protein Rodent Diet (Envigo, Madison, Wisconsin USA), chow commonly used by research institutions (see e.g. [29]). To confirm that *Salmonella* does not lose viability on chow, pieces of chow (approximately 5 mm in diameter) were prepared from pellets (Fig 1A) and inoculated with known numbers of bacteria. Following incubation at room temperature for 1 or 3 h, the chow pieces were homogenized, diluted and plated to enumerate colony forming units (CFUs). As a comparison, bacteria were also diluted and inoculated in sterile pharmaceutical grade saline (SPGS). *Salmonella* showed no decrease in viability in food or saline over the course of 3 h (Figs 1B and C), irrespective of the initial numbers of bacteria.

**Fig 1:**
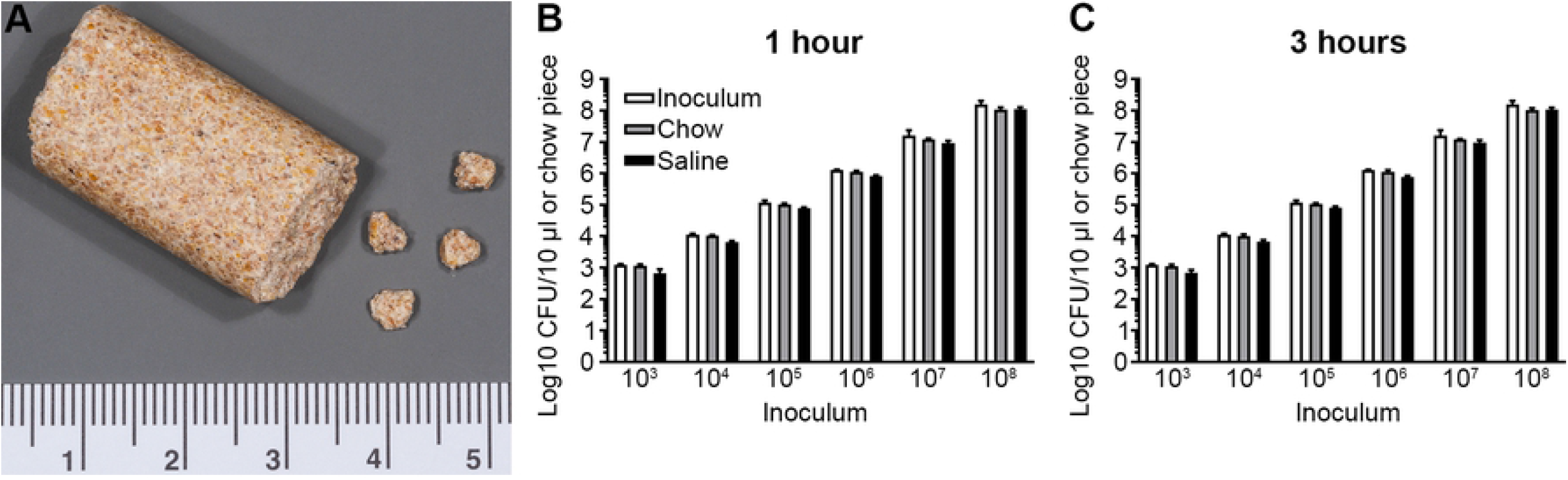
Salmonella survives on pieces of mouse chow. (A) Representative image of structure and size of a regular mouse chow pellet and prepared chow pieces. (B, C) Survival of *Salmonella* on pieces of chow and in SPGS, 1 and 3 h after inoculation. Original inoculum added as a comparison. Data represent the mean ± standard deviation of three independent experiments (n=1 per experiment).

### Delivery of *Salmonella* using chow results in infection similar to gavage delivery and has less variability

Gavage is the method traditionally used to deliver *Salmonella* in oral infection of mice and for streptomycin treatment. To investigate how oral infection with chow compared to the commonly used procedure for infection by gavage, streptomycin treated (hereafter referred to as strep+) C57Bl/6J mice were infected with 10^4^ CFU *Salmonella* using either method (Fig 2A). For oral gavage, mice were gavaged with streptomycin (20 mg) 24 h prior to being gavaged with *Salmonella* diluted in SPGS (100 μl). Mice were fasted for 4 h prior to each gavage. For mice infected with chow, to avoid gavage altogether, streptomycin was added to the drinking water (final dilution of 5 mg/ml) for 24 h [30]. C57Bl/6J mice drink approximately 6 ml of water per 24 h [31], which results in an approximate total dose of 30 mg streptomycin. After 24 h normal drinking water was returned, mice were fasted for approximately 20 h, then fed pieces of chow inoculated with *Salmonella* diluted in SPGS. Since we were concerned that high levels of *Salmonella* might affect the palatability of chow, we initially used a low inoculum of *Salmonella*, although the dose most frequently used is approximately 10^8^ *Salmonella* [16, 29, 32]. Mice readily ate chow pieces inoculated with approximately 10^4^ CFU. At 3 days post infection (p.i.), mice displayed very mild, if any, clinical signs of disease, although feces was frequently found on the walls of the cage, indicating wet stool. Despite the low inoculum, all mice were infected although those infected with inoculated chow had higher bacterial loads in tissues, especially in the intestines, compared to those infected by gavage (Fig 2B). While this could be a result of the inoculation method itself, it could also be due to either the prolonged streptomycin treatment in the drinking water or the prolonged fasting period [33, 34]. Bacterial numbers were more consistent between the mice in the chow infection group when compared to mice in the gavage group. This consistency was most notable in the feces, where bacterial numbers in gavaged mice ranged from 0 (below the limit of detection) to 3.4×10^8^ and in chow infected mice from 1.6×10^8^ to 1.3×10^10^. The variability in infection due to oral gavage has been observed previously [26]. Since oral gavage may induce regurgitation and subsequent unintended tracheal delivery of substances [22] we also examined the lungs for *Salmonella*.Mice infected by either method contained low levels of *Salmonella* in the lungs, again with more variability in mice infected with gavage (0 to 8.7×10^3^) compared to those infected via chow (8.9×10^1^ to 5.4×10^2^) (Fig 2B). The presence of bacteria in the lungs is not overly surprising considering the susceptibility of strep+ C57Bl/6J mice to disseminated *Salmonella* infection, however, two mice infected by oral gavage contained considerably (>1 log) higher bacterial numbers in the lungs than the rest of the mice. This may indicate unintended tracheal administration of *Salmonella* as a complication of the gavage process. Altogether, these findings indicate that infection with inoculated chow results in more consistent disease with less complications compared to the traditional gavage method.

**Fig 2:**
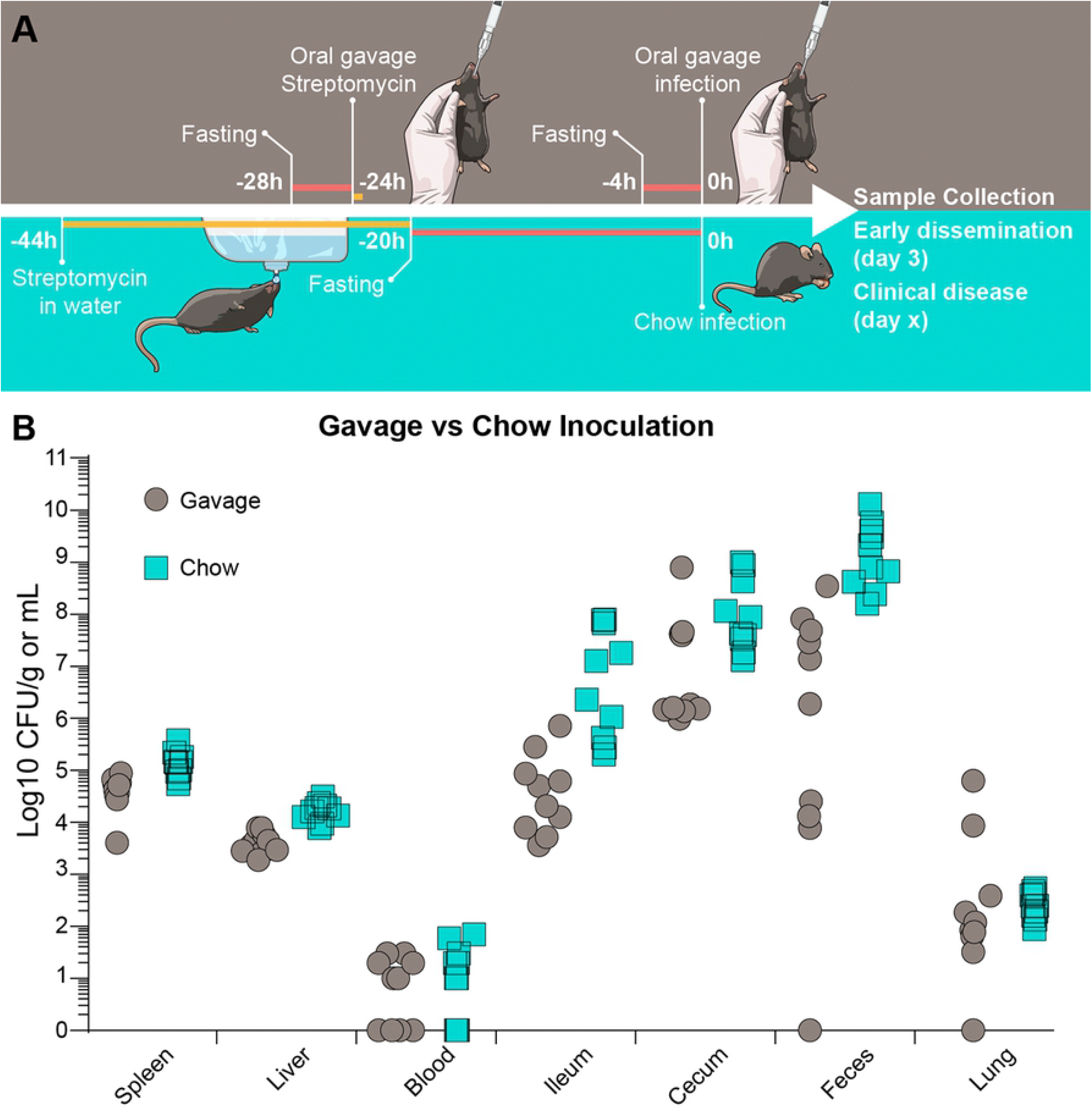
Infection via inoculated chow or oral gavage results in similar disease progression. (A) Schematic representation of the gavage and chow infections performed in this study. (B) Bacterial numbers in tissues from mice infected either by oral gavage or pieces of chow 3 days p.i. n=10 mice. Symbols represent individual mice.

### Mice readily eat chow pieces inoculated with high loads of *Salmonella* and show consistent infection

Having shown that chow inoculated with a low dose (10^4^ CFU) of *Salmonella* results in consistent infection we next compared a dose range. Pieces of chow were inoculated with 10^3^ to 10^6^ CFU and fed to strep+ C57Bl/6J mice. For doses of 10^5^ bacteria and higher, the bacteria were pelleted by centrifugation and then diluted in SPGS (washed) before dilution since mice were hesitant to eat chow inoculated with these high doses if they were not washed (data not shown). This may be due to the intrinsic smell or taste of the growth medium or the presence of high amounts of bacterial products. After introducing the wash step, mice readily ate chow pieces containing up to 10^8^ CFU. By 3 days p.i. all mice had developed systemic infection even at the lowest dose (10^3^) and bacterial loads were strikingly similar (Fig 3A), indicating no dose dependence for the development of disseminated systemic disease, correlating with previous reports [30]. Bacterial loads were consistent in systemic tissues but showed more variability in gastrointestinal tissues with the greatest variability seen in the ileum (3.0×10^5^ to 5×10^8^ CFU/g). Bacterial loads were highest in the cecum (approximately 10^9^ CFU/g) and lowest in the blood (approximately 10^2^ CFU/ml). To investigate if chow infection works in a resistant mouse strain, we used C57Bl/6J mice reconstituted with a functional Nramp1 (hereafter referred to as Nramp1^+/+^) [11, 13, 14]. Mice with a functional Nramp1 protein are more resistant to oral *Salmonella* infection both with and without prior streptomycin treatment [8, 17]. For the Nramp1^+/+^ mice we tested doses of 10^4^ and 10^5^ bacteria. A dose of 10^4^ resulted in inconsistent infection in systemic tissues, while 10^5^ resulted in a consistent systemic infection, indicating that 10^5^ is the minimum dose required for 100% systemic infection. As expected, Nramp1^+/+^ mice were substantially more resistant to infection, compared to C57Bl/6J mice, with lower bacterial loads in all tissues but particularly in systemic organs (Fig 3B). Nonetheless, bacterial loads in Nramp1^+/+^ mice followed the same trends as in C57Bl/6J mice with the highest number of bacteria in the feces followed by the cecum, ileum, spleen, liver and blood. Interestingly, bacterial loads were also lower in the feces of Nramp1^+/+^ mice, indicating that functional Nramp1 is important for limiting bacterial numbers in intestinal luminal contents as well as systemically.

**Fig 3:**
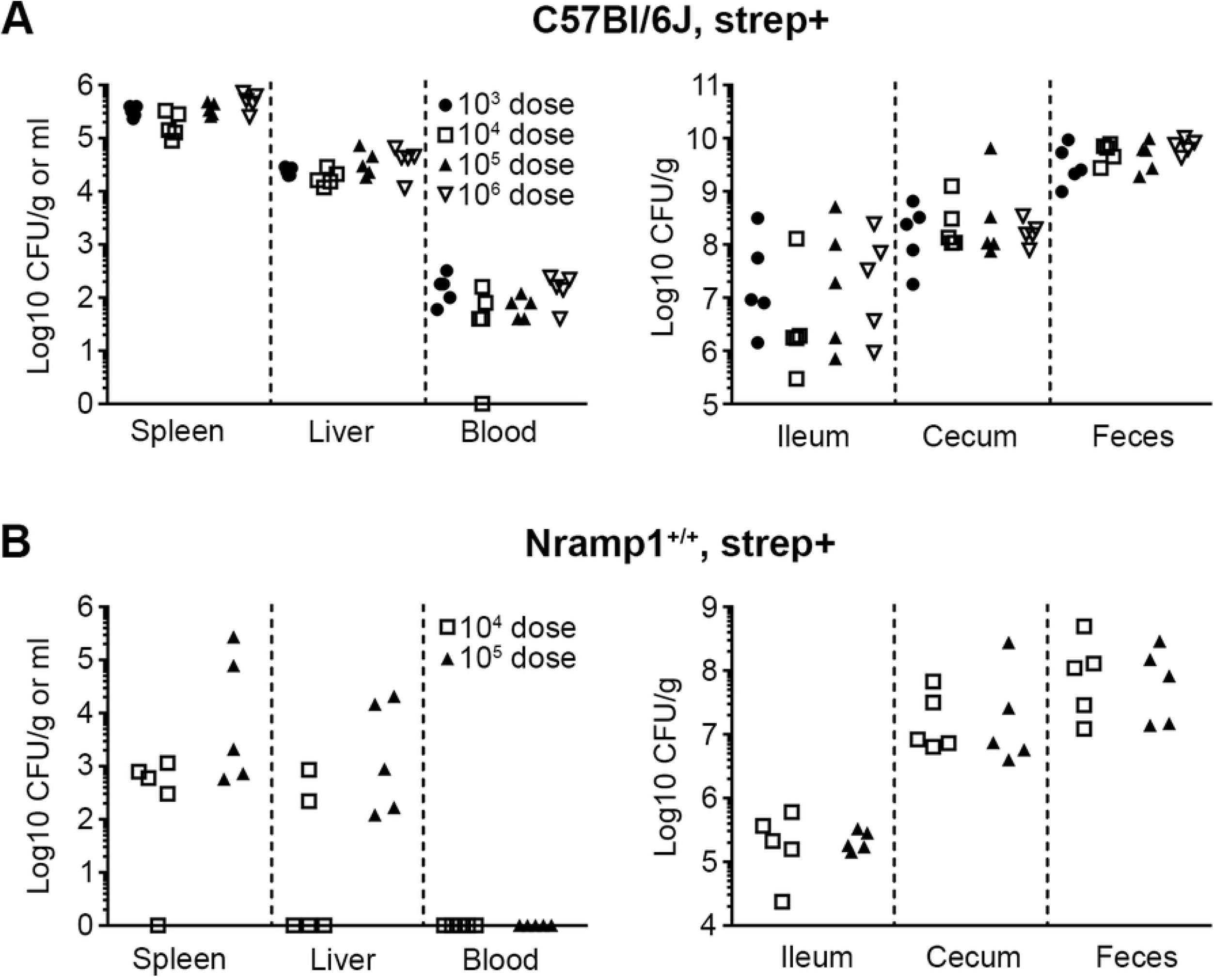
Pieces of chow containing a wide range of inoculums can be fed to strep+ mice. (A) Bacterial numbers in tissues 3 days p.i. in strep+ C57Bl/6J mice infected with different doses. (B) Bacterial numbers in tissues 3 days p.i. in strep+ Nramp1^+/+^ mice infected with different doses. n=5 mice and symbols represent individual mice. Tissues where bacterial load was below the level of detection are given a value of “1” for visualization purposes.

Next we infected non-streptomycin treated (hereafter referred to as strep-) C57Bl/6J and Nramp1^+/+^ mice using pieces of chow. Since these mice are much less susceptible to oral infection, we used a dose of 10^8^ bacteria, the standard dose used in oral infections of mice with *Salmonella* (see e.g. [16, 32, 35]). After washing the bacteria, mice readily ate pieces of chow containing this high inoculum. C57Bl/6J mice had consistent bacterial loads in intestinal tissues, while Nramp1^+/+^ mice showed more variability in loads which were also sometimes below the level of detection (Fig 4). In systemic tissues, no bacteria were detected in any of the Nramp1^+/+^ mice and bacterial numbers were variable in C57Bl/6J mice. In the intestinal tract, the differences in bacterial loads between C57Bl/6J and Nramp1^+/+^ mice were statistically significant in the ileum and feces.

**Fig 4:**
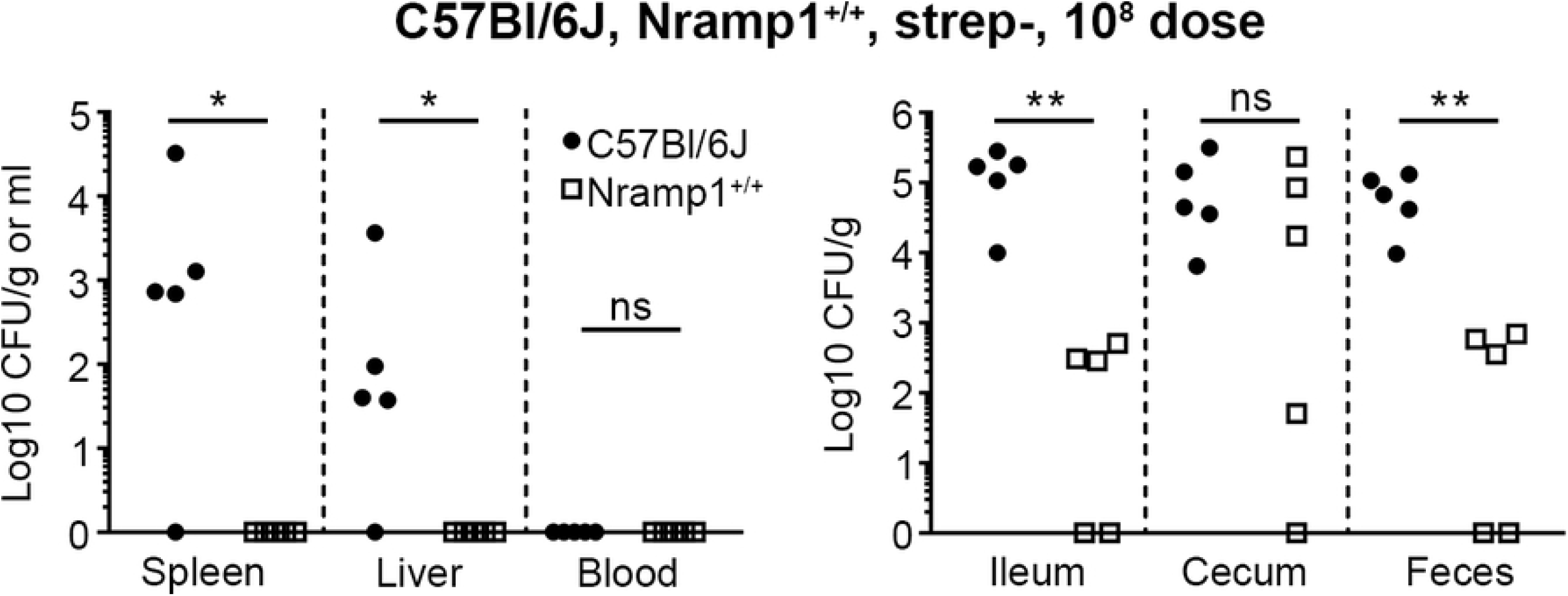
Infection of strep- mice results in more variability in organ loads of *Salmonella*. Bacterial numbers in tissues 3 days p.i. in strep- C57Bl/6J and Nramp1^+/+^ mice infected with 10^8^ CFU *Salmonella*. n=5 mice and symbols represent individual mice. Tissues where bacterial load was below the level of detection are given a value of “1” for visualization purposes. Asterix indicates statistical significance; * p < 0.05, ** p < 0.01, ns, not statistically different, two-tailed Mann-Whitney U test.

In summary, infections of strep+ and strep- C57Bl/6J and Nramp1^+/+^ mice with pieces of chow inoculated with *Salmonella* are consistent with studies using oral gavage. While it is difficult to determine the dose of nontyphoidal *Salmonella* required to cause gastroenteritis in humans, reports indicate doses as low as 10 organisms [36, 37]. In this infection model we achieved consistent infection of strep+ mice using only 10^3^ organisms, indicating that this model can be used to investigate a range of relevant infection doses.

### Mice infected using pieces of chow succumb to *Salmonella* infection similar to mice infected by oral gavage

Next, we sought to compare the development of clinical disease of C57Bl/6J and Nramp1^+/+^ mice infected using inoculated chow. Strep+ and strep- mice were inoculated with 10^4^ or 10^8^ CFU of *Salmonella*, respectively, and monitored for overt clinical signs at which point they were euthanized. As expected, strep+ mice succumbed faster to disease compared to strep- mice (Figs 5A and B). Only one strep- Nramp1^+/+^ mouse developed clinical signs and was euthanized 15 days p.i.; all others were euthanized at the experimental end point of 21 days p.i. This is consistent with an intact microbiota and the ability of Nramp1^+/+^ mice to effectively combat infection. The mouse euthanized 15 days p.i. showed very high bacterial loads in all tissues except the liver and blood (open symbols in Fig 5F). Strep+ mice developed very high bacterial loads in the intestinal tissues, especially C57Bl/6J mice, which also developed high loads in systemic tissues (Figs 5C and D). This indicates that Nramp1 contributes to controlling *Salmonella* infection both in systemic and intestinal sites. At the time of euthanasia, strep- C57Bl/6J mice had higher and more consistent bacterial loads in all tissues compared to strep- Nramp1^+/+^ (Figs 5E and F). It is interesting to note that although strep- C57Bl/6J mice have lower intestinal bacterial loads compared to strep+ mice, they do go on to develop higher systemic bacterial loads. This is probably a result of the prolonged survival of strep- mice allowing additional time for the bacteria to replicate at systemic sites. These findings correlate with other studies of oral infection of strep- susceptible and resistant mouse strains [8, 35, 38], and show the reduced survival of strep+ mice.

**Fig 5:**
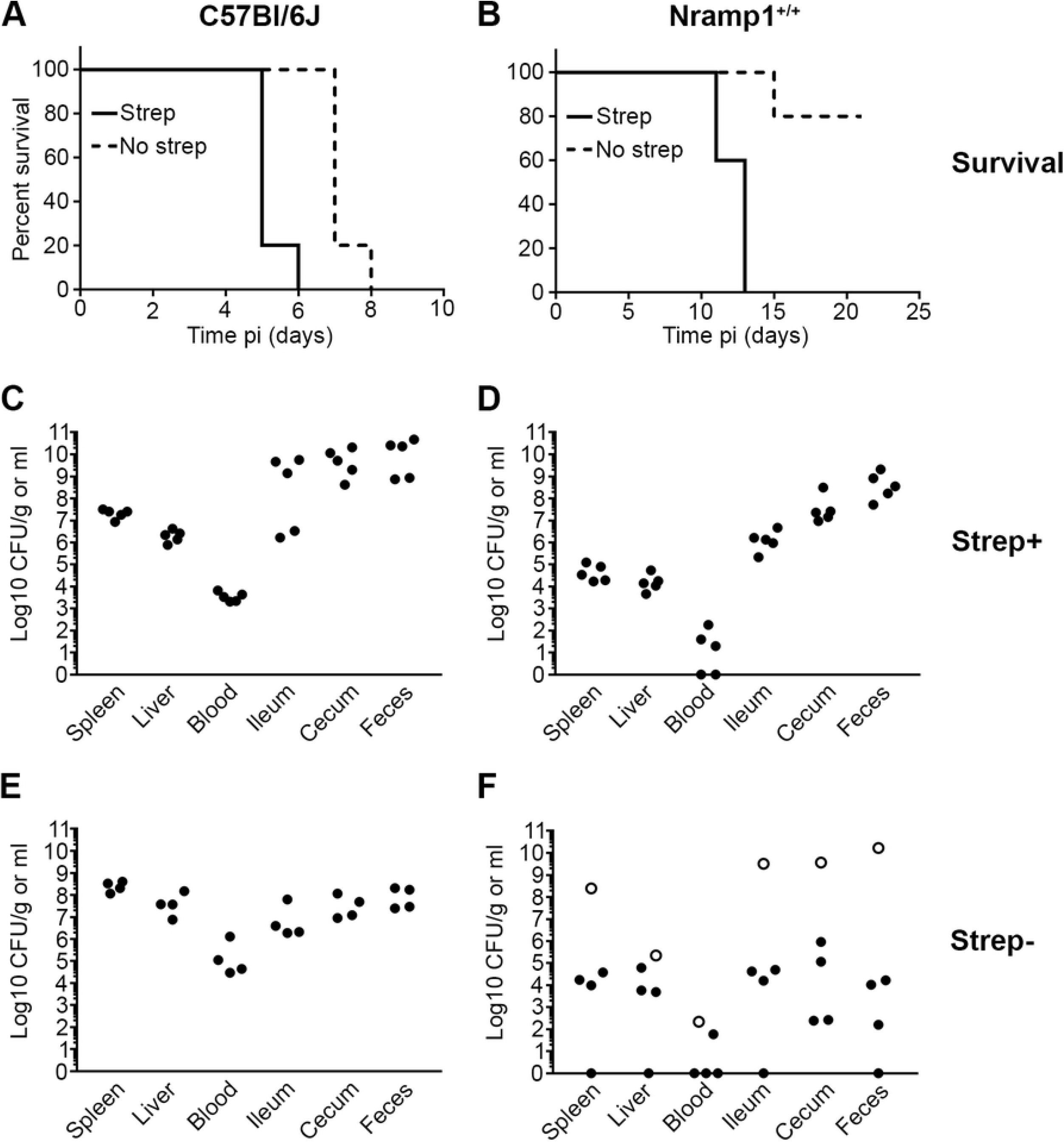
Strep- mice survive longer than strep+ mice following chow infection and show more variability in organ loads. (A) Survival of strep+ and strep- C57Bl/6J mice orally infected with 10^4^ and 10^8^ bacteria, respectively. (B) Survival of strep+ and strep- Nramp1^+/+^ mice orally infected with 10^4^ and 10^8^ bacteria, respectively. (C) Bacterial loads of strep+ C57Bl/6J mice in (A) at time of euthanasia. (D) Bacterial loads of strep+ Nramp1^+/+^ mice in (B) at time of euthanasia. (E) Bacterial loads of strep- C57Bl/6J mice in (A) at time of euthanasia. (F) Bacterial loads of strep- Nramp1^+/+^ mice in (B) at time of euthanasia. n=5 mice in each experiment and symbols represent individual mice. One strep- C57Bl/6J mouse died, leaving only 4 mice for tissue collection (Fig 5E). Open symbols in Fig 5F indicate the one Nramp1^+/+^ mouse that was euthanized 15 days p.i. Tissues where bacterial load was below the level of detection are given a value of “1” for visualization purposes.

### Shorter periods of fasting increases hesitancy of mice to eat inoculated chow

For practical reasons, it would be advantageous to have the option of varying the fasting time although mice fasted for shorter times are less likely to eat inoculated chow. Therefore, to determine how changing the duration of fasting would impact the method, we fasted mice for 4, 8 or 14 h and then measured the time taken to completely consume inoculated pieces of chow. These mice were not streptomycin treated. After 14 h (o/n) of fasting, mice consumed chow within 2 min, while shorter fasting resulted in longer consumption times, up to 23 min (Fig 6A). The inoculum did not seem to affect consumption time, since mice fasted for 4 h and offered chow inoculated with 10^8^ CFU consumed the chow in a time frame similar to mice fed 10^4^ CFU (Fig 6A). The very low consumption time after 14 h may be due to fasting taking place overnight when mice are more active. Shorter fasting times required a modification to the feeding procedure, where mice were moved into individual clean cages, offered a piece of chow and then left undisturbed until consuming the whole chow piece. When comparing the organ loads 3 days p.i. of mice fed 10^8^ CFU after fasting for 4 h (Fig 6B) or 20 h (Fig 4), the shorter fasting period led to lower bacterial loads in systemic tissues and in some of the intestinal tissues. This indicates that shorter fasting times results in a delayed, or less efficient, dissemination of *Salmonella*. In a study describing oral infection of mice with *Listeria monocytogenes* the authors noted that mice sometimes had to be left undisturbed for up to 2 h to eat the offered food, even after 24 h of fasting [28]. The reason for the mice more readily eating the inoculum in this study may be due to the inoculated pieces of chow are derived from their regular chow, a type of food they are already familiar with.

**Fig 6:**
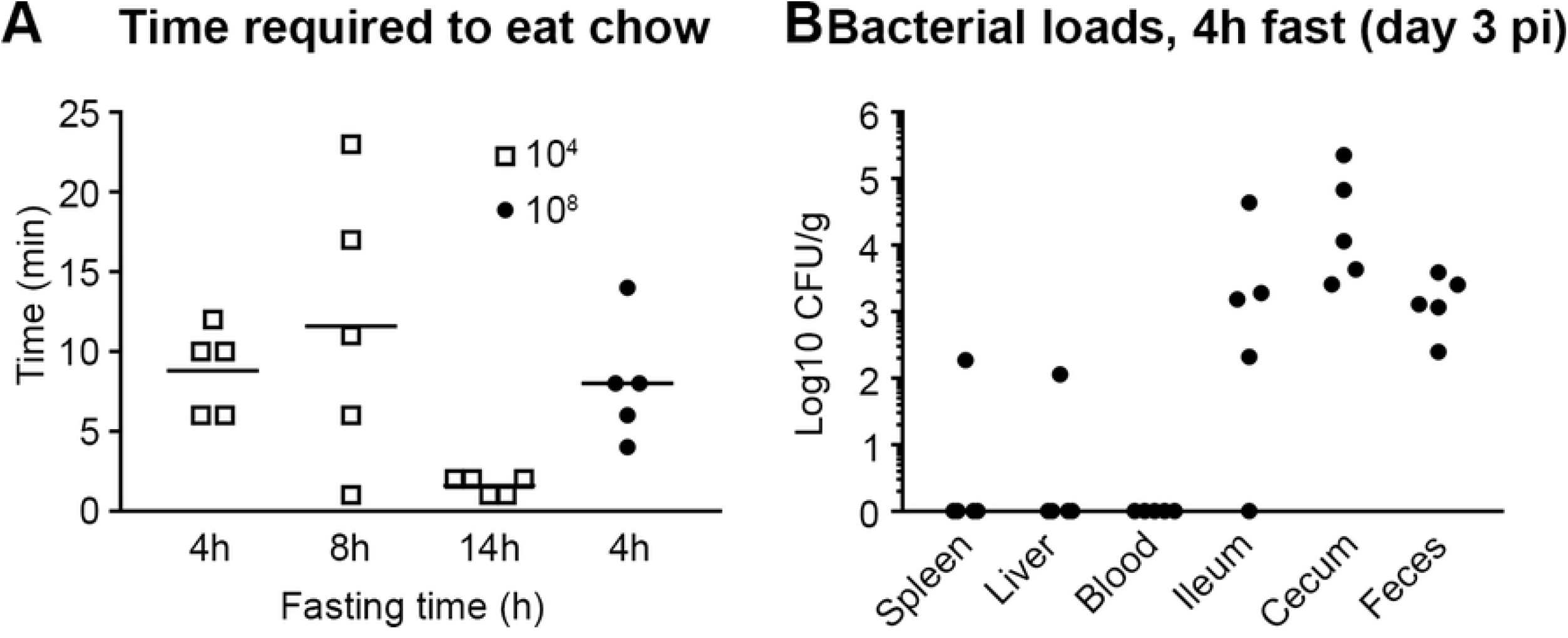
Fasting time affects the time taken to eat inoculated chow and the progression of infection. (A) Time required for mice to completely consume pieces of chow with the indicated inoculum and fasting time. Symbols represent individual mice and horizonatal line represents the mean. 5 mice per experiment. (B) Bacterial loads in tissues 3 days p.i. following 4 h of fasting and an inoculum of 10^8^ CFU *Salmonella*. n=5 mice, symbols represent individual mice.

In summary, oral infection of mice using pieces of their regular chow inoculated with *Salmonella* mimics the natural route of infection, is technically very simple and results in reproducible infection while avoiding the stress and potential adverse side effects of oral gavage. Further refinements to the method are possible, such as; adjusting fasting times; the concentration of streptomycin in drinking water; and the time allowed for mice to access water containing streptomycin. We expect that this approach would work for other strains of mice and intestinal pathogens.

## Materials and Methods

### Ethics statement

All animal studies were carried out following the recommendations in the Guide for the Care and Use of Laboratory Animals, 8^th^ Edition (National Research Council), and the animal study protocol was approved by the Rocky Mountain Laboratories Animal Care and Use Committee. Protocol number 2017-021-E. Animals were either euthanized before the development of clinical disease, at specified time points, or at the defined humane endpoint (development of clinical disease: ruffled fur, hunched posture, lethargy).

### Bacterial strains and growth conditions

*Salmonella* Typhimurium strain SL1344 was used for all experiments. For infections, bacteria were grown in a 125 ml Erlenmeyer flask in 10 ml LB-Miller containing 100 μg/ml streptomycin for 18 h at 37°C, with shaking at 225 RPM, and diluted in sterile SPGS to get the correct inoculum in 10 μl (e.g. to get 10^4^ CFU an overnight culture was diluted 1:5000), the volume added to pieces of chow. For inoculum of 10^5^ CFU and higher, a wash step was included prior to dilution. 1 ml of the overnight culture was centrifuged at 8000 G for 2 min, the supernatant was aspirated, and the bacterial pellet resuspended in 1 ml SPGS. In order to achieve a final concentration of 10^8^ CFU in 10 μl, the bacterial pellet was resuspended in 0.5 ml SPGS.

### Preparation of chow pieces for infection

Mouse chow pellets (2016 Teklad Global 16% Protein Rodent Diet, Envigo, Madison, Wisconsin USA), were broken into smaller pieces of about 4-5 mm in diameter by gentle tapping with a small hammer followed by trimming with forceps. Selected pieces were gently tested for physical integrity, by dropping from a height of 4-5 inches, before 10 μl of inoculum was pipetted onto the surface. Prepared pieces were kept separated in a petri dish during transport to the animal facility. One piece of inoculated chow was retained for estimation of the CFU by plating.

### Mouse chow infections

Except where specified, mice had unlimited access to food and water. For Streptomycin pretreatment the antibiotic (5 mg/ml) was added to drinking water 42 – 46 h prior to infection for 24 h. Mice were then moved to a clean cage (to limit coprophagy and access to cached food), containing normal drinking water but no chow. After a period of 18 – 22 h (typically 20) individual mice were put in a clean empty cage (without bedding material) and were then offered a piece of chow. Typically, mice ate the piece of chow immediately or within a couple of min. For short fasting times (4 and 8 h) mice were left undisturbed until the chow was eaten. Immediately after the inoculated chow was consumed, mice were returned to their cage with unlimited access to food and water.

### Mouse oral gavage infections

Mice were streptomycin treated 24 h before infection, using a blunt end gavage needle with 100 μl SPGS containing 200 mg/ml streptomycin. For *Salmonella* infection, mice were gavaged with bacteria in 100 μl SPGS. Mice were fasted for 4 h prior to all gavages. For infections without streptomycin treatment, mice were only fasted prior to feeding.

### Tissue collection and processing

Mice were euthanized by isoflurane inhalation followed by exsanguination. Tissues were collected in screwcap tubes containing 500 μl SPGS and 3-4 2.0 mm zirconia beads (BioSpec Products) and homogenized using a Bead Mill 24 (Fisher Scientific, 4.85 m/s for 20 seconds). Tubes were weighed before and after organ collection. CFUs were estimated by 10 μl spot plating of 10-fold dilutions on LB agar plates containing the appropriate antibiotic.

## Acknowledgments

We thank the members of the Steele-Mortimer laboratory, Karin Peterson and Clayton Winkler for critical review of the manuscript, and Ryan Kissinger for assistance with figures. This research was supported by the Intramural Research Program of the NIH, NIAID.

